# A Thermodynamic Framework Linking Black Box Growth models with Genome Scale Metabolite models

**DOI:** 10.64898/2026.09.14.751451

**Authors:** Pujiang Jia, Deyu Wang, Hualin Shi

## Abstract

Genome scale metabolic models provide detailed mechanistic descriptions of cellular metabolism, whereas black box growth models capture physiological behaviors using a small number of effective parameters. However, the quantitative relationship between these two modeling scales remains unclear. In this study, we develop a thermodynamic framework that connects black box growth models with thermodynamically constrained genome scale metabolic models. By linking black box model parameters with Gibbs energy dissipation rates derived from genome scale metabolic models, we demonstrate that coarse grained physiological descriptions can be obtained from detailed metabolic networks while preserving their thermodynamic foundation. We validate this framework in both Escherichia coli and yeast, showing that the resulting black box models reproduce key physiological behaviors, including growth dynamics, biomass yield, and overflow metabolism observed experimentally. Our results indicate that black box growth models and genome scale metabolic models are connected through shared thermodynamic constraints, revealing a consistent thermodynamic basis across different levels of metabolic description. This framework provides a general approach for integrating detailed metabolic networks with simple black box growth models and enables efficient multiscale modeling of cellular metabolism.

## Introduction

Life is one of the most remarkable manifestations of non equilibrium thermodynamics. The ability of living systems to grow and reproduce distinguishes them fundamentally from inanimate matter, yet this extraordinary complexity does not exempt them from the universal laws of physics.The pursuit of understanding how these physical principles govern biological growth led to the development of quantitative descriptions of microbial growth in the second half of the twentieth century [1]. The pioneering studies of Jacques Monod [2, 3], which introduced biomass yield concepts and the Monod growth law, laid the foundation for quantitative microbial growth modeling. The development of continuous culture systems by Herbert and colleagues [4] in 1956 provided a powerful experimental framework for investigating microbial kinetics under controlled conditions. Subsequently, Herbert [5] and Pirt [6] independently introduced maintenance concepts to explain the variation of biomass yield with dilution rate. Thermodynamic considerations were further incorporated into microbial growth models through the introduction of the degree of reduction (*γ*) [7, 8]. Minkevich and Eroshin [8] derived energetic relationship linking *γ* to biomass yield and established upper thermodynamic bounds as expressed by Thornton’s rule [9]. Roels [10, 11] reformulated microbial yield limits to ensure consistency with the second law of thermodynamics for open systems. More important, Battley [12–14] developed methods to estimate the Gibbs free energy and enthalpy of formation of biomass.With these theories, researchers adopted black box model [15–17] based on overall mass and energy balances. In this approach, the cell was treated as a system that converts substrates into biomass and byproducts according to an overall macrochemical reaction [18].black box model relied on measurable input output fluxes in bioreactors and on global conservation principles without resolving intracellular pathway structure [10, 11, 19–22]. Despite the coarse grained representation, it successfully related biomass yields to Gibbs free energy dissipation and provided general energetic limits for microbial growth [15–17]. Recent developments in coarse grained black box models have demonstrated their potential for capturing bacterial resource allocation strategies and linking molecular mechanisms with system level physiological behaviors [23, 24].

The emergence of high throughput technologies has changed the level at which cellular metabolism can now be analyzed. Genome scale metabolic models [25] describe metabolism at the level of individual reactions and enable the quantitative analysis of intracellular flux states under various constraints, such as enzyme constrains [26], thermodynamic cosntrains [27], dFBA [28] and SRFBA [29]. Various works in solution space sampling [30–32] had been made to explain and predict growth rate fluctuations and universal scaling relation.

Previous studies [27, 33, 34] have established important conceptual links between black box model and genome scale metabolic model by incorporating thermodynamic principles into genome scale model and extending black box formulations to describe genome scale simulation results. However, a quantitative understanding of how the high dimensional information contained in genome scale flux solutions can be captured by a low dimensional black box representation remains lacking.

In this study,a genome scale thermodynamic constrained metabolic model of *Escherichia coli* is used to calculate the key energetic parameters of the black box model framework, thereby establishing a quantitative link between these two levels of description.By bridging classical energetic theories and modern genome scale modeling, this work contributes to a deeper theoretical understanding of microbial growth across different hierarchical levels.We focus on metabolic networks from *Escherichia coli*, and our approach is applicable to studying the metabolism and growth of other bacteria.

## Black box model

### Herbert–Pirt substrate distribution relation

The black box model neglects detailed intracellular metabolic network and instead describes cellular metabolism in terms of nutrient uptake, biomass formation, and the secretion of metabolic products. By imposing overall mass and energy conservation constraints, it provides a compact representation of cellular metabolism that typically depends on only a small number of macroscopic parameters. The model is particularly useful for the design and analysis of batch, continuous, and fed batch processes, whose characteristic timescales are typically on the order of hours. Within this framework, the Herbert–Pirt equation describes how substrate uptake is partitioned among biomass formation, anabolic product formation, and cellular maintenance [5, 6]:

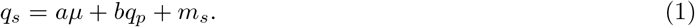

In this formulation, substrate intake(at rate *q*_*s*_) is consumed for biomass formation (at growth rate *µ*),anabolic products(at rate *q*_*p*_), excluding fermentation byproducts such as acetate and ethanol, and maintenance(*m*_*s*_). The parameter 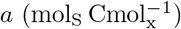 denotes the substrate required for biomass formation, 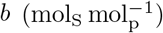 denotes the substrate required for anabolic product formation, and *m*_*s*_ denotes the substrate consumption associated with maintenance. During cellular metabolism, the Gibbs free energy released by biochemical reactions is ultimately dissipated. A substantial fraction of this energy is converted into heat and transferred to the surrounding environment, thereby contributing to the thermal conditions under which temperature dependent cellular processes occur. When heat removal is limited, metabolic heat accumulation may alter the local temperature and consequently affect microbial growth. In addition, a portion of the dissipated Gibbs energy is required for cellular maintenance, including the turnover and repair of cellular components, the preservation of membrane gradients, and other essential processes that do not directly contribute to biomass formation. The total energy dissipation is:

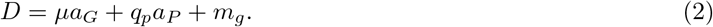

*a*_*G*_ and *a*_*P*_ represent the Gibbs free energy dissipation requirements associated with the formation of one C-mol of biomass and anabolic product, respectively, during heterotrophic growth, while the maintenance Gibbs energy requirement, *m*_*g*_, is assumed to primarily depend on temperature because the underlying maintenance processes, such as cellular component turnover and restoration of membrane gradients, are governed by temperature dependent biochemical reaction rates. Based on experimental data from diverse microorganisms, Tijhuis [35] observed an Arrhenius type dependence of *m*_*g*_, leading to the empirical correlation:

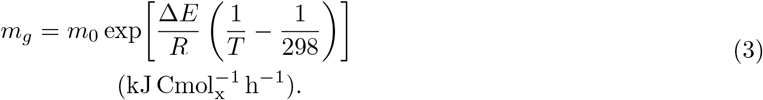

Here, the exponential term accounts for the temperature dependence of maintenance energy, while *m*_0_ represents the fitted value of *m*_*g*_ at the reference temperature of 298 K. Under aerobic and anaerobic conditions, the basal maintenance energy(*m*_0_) are estimated to be 5.7 and 3.3 kJ 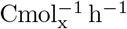, respectively. The activation energy (Δ*E*) associated with maintenance processes remains comparable between the two systems, with a value of approximately 6.9 × 10^4^ J mol^−1^. To calculate *m*_*s*_, we use the catabolic Gibbs free energy of the substrate under standard conditions (Δ_cat_*G*(*standard*) in Table 1), such that *m*_*s*_ = *m*_*g*_*/*Δ_cat_*G*. The Gibbs free energy dissipation requirement for heterotrophic growth, denoted by 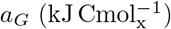, was estimated using the empirical correlation proposed by Heijnen, which was parameterized using carbon sources spanning approximately 1 ≤ *c* ≤ 6 and 0 ≤ *γ* ≤ 8 [17]:

**Table 1.**
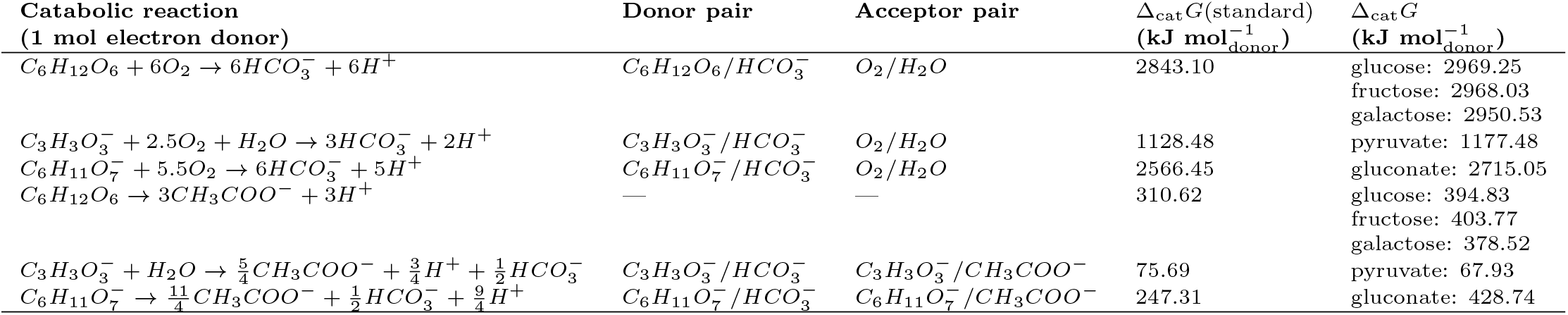
Catabolic reactions for one mole of electron donor and corresponding Gibbs free energy changes. Δ_cat_*G*(standard) values are standard Gibbs energy changes of catabolic reactions for the specific carbon sources at *pH* = 7. Δ_cat_*G* values are Gibbs energy changes derived from thermoFBA, including logarithmic concentration corrections.

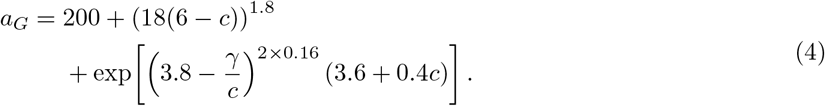

In this correlation, *c* represents the carbon chain length of the substrate and *γ* denotes its degree of reduction. The carbon chain contribution describes the effect of substrate structure on Gibbs energy dissipation, with decreasing dissipation as *c* approaches 6 within the carbon source set considered by Heijnen [17]. The degree of reduction reflects the redox state of the substrate. A lower dissipation is generally observed for substrates with *γ* ≈ 4, which is close to the reduction degree of biomass (*γ*_*X*_ ≈ 4.2 when 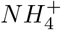 serves as the nitrogen source). Thus, substrates with redox characteristics closer to biomass formation require less dissipated Gibbs energy.

The parameter 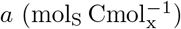 is a central quantity in the black box model. During biomass formation, part of the substrate is used for anabolism and the remainder is oxidized to supply the required Gibbs free energy:

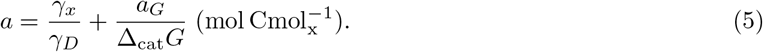

Here, *γ*_*x*_ and *γ*_*D*_ denote the degrees of reduction of biomass and substrate per C atoms, respectively. The first term, *γ*_*x*_*/γ*_*D*_, reflects the constraint imposed by the transfer of reducing equivalents during biomass synthesis. In contrast, *a*_*G*_ represents the Gibbs free energy requirement for heterotrophic growth, while Δ_cat_*G* corresponds to the Gibbs free energy released from substrate catabolism. Therefore, the second term, *a*_*G*_*/*Δ_cat_*G*, captures the thermodynamic conservation associated with the energetic cost of biomass formation.

If the supplied carbon source is assumed to be converted only into biomass and catabolism byproducts(*CO*_2_, acetate, ethanol etc.), the anabolic byproduct term vanishes and *bq*_*p*_ = 0. The biomass yield, *Y*_*x*_, is defined as the mass of biomass formed per unit mass of substrate consumed. In terms of the specific growth rate and substrate uptake rate, this definition can be written as:

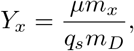

where *m*_*x*_ denotes the molar mass of biomass and *m*_*D*_ denotes the molar mass of the substrate. Under the no byproduct assumption, the two remaining contributions to Eq. (1) are the growth associated term *aµ* and the maintenance associated substrate consumption *m*_*g*_*/*Δ_cat_*G*. Substituting Eq. (5) for *a* therefore gives:

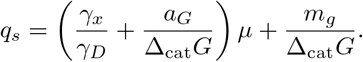

Substituting this *q*_*s*_ relation into the yield definition above, yields:

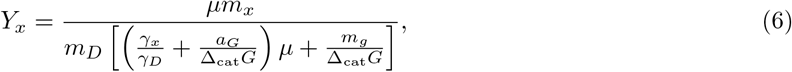

which gives the biomass yield as an explicit function of the specific thermodynamic parameters(*a*_*G*_, Δ_cat_*G*),substrate dependent parameters(*m*_*D*_, *γ*_*D*_) and growth rate(*µ*).

### Genome scale model

Flux balance analysis (FBA) is a widely used framework for analyzing biochemical networks, particularly genome scale metabolic reconstructions [25, 36–38]. In this study, we used a thermodynamic constrained genome scale model based on the *E. coli* iJR904 reconstruction [27]. This model is particularly suitable for the present analysis because it explicitly accounts for Gibbs free energy dissipation and therefore enables direct analysis of energetic constraints on metabolism.

The model contains 626 unique metabolites and 917 metabolic reactions, including 724 biochemical reactions, 193 transport reactions, and 144 exchange reactions. In addition to standard mass balance constraints, it includes proton balance, charge balance, and Gibbs energy balance constraints. All reactions are reversible in principle, but their feasible directions are restricted by the second law of thermodynamics (see Appendix).

Another thermodynamically constrained genome scale model of *S. cerevisiae*, developed based on genome scale metabolic model iND750. Following the same thermodynamic formulation as the *E. coli* model, this model integrates stoichiometric and thermodynamic constraints. The resulting model contains 156 metabolites, 241 metabolic processes, and 15 exchange processes.

## Establishing a thermodynamic bridge between black box models and genome scale models

### Extended black box model

Conventional black box model cannot reproduce overflow metabolism during bacterial growth: a metabolic strategy widely observed across both prokaryotes and eukaryotes. Overflow metabolism typically emerges when the carbon source is abundant, even under fully aerobic conditions, cells divert part of their carbon flux into fermentative pathways, sacrificing biomass yield in exchange for a higher rate of energy generation that sustains rapid growth.To capture this behavior, we extend the conventional black box model based on the framework proposed by [34] as follows. We assume two metabolic modes.The first is respiration, which completely oxidizes the carbon source to *CO*_2_ and *H*_2_*O* and releases a larger amount of Gibbs free energy, Δ_cat_*G*_2_ 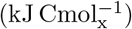, as listed in Table 1. This mode contributes a growth rate component *µ*_2_, and its Gibbs free energy requirement *a*_*G*2_ 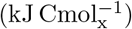 is approximated from Heijnen’s relation in (5). The second is fermentation, which converts the carbon source into byproducts such as acetate and releases less Gibbs free energy, Δ_cat_*G*_1_ 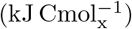. This mode contributes a growth rate component *µ*_1_, and its Gibbs free energy requirement *a*_*G*1_ 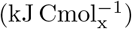 is assumed to be 50–150 kJ 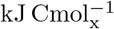 lower than *a*_*G*2_ [34]; here we use *a*_*G*1_ = *a*_*G*2_ − 120. The total growth rate is then:

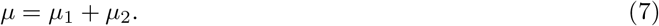

The total energy dissipation rate reads:

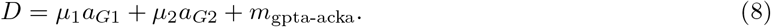

*m*_gpta-acka_ is a piecewise quantity: under growth before overflow it reduces to *m*_*g*_, whereas under overflow metabolism it denotes the maintenance energy specific to the overflow regime, which differs from that of either the pure fermentation or the pure respiration pathway. Accordingly, the Herbert-Pirt relation in Eq. (1) becomes:

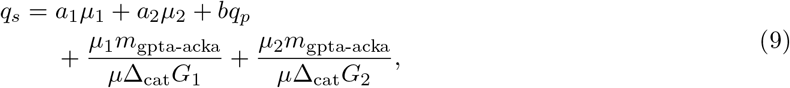

where *a*_1_ and *a*_2_ denote substrate required for biomass formation of fermentation and respiration pathways respectively.In our model, both fermentation and respiration pathways don’t produce byproduct except for catabolism byproducts(*CO*_2_,acetate,ethanol etc.), so the anabolic byproduct term vanishes and *bq*_*p*_ = 0.

According to Niebel [27], bacterial growth is subject to an upper limit on the Gibbs free energy dissipation rate(*D*_max_):

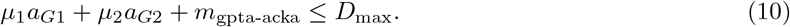

Using the value reported in [27], *D*_max_ = 4.9 kJ gDW^−1^ h^−1^, and the biomass composition used in our model, *CH*_2.3544_*O*_0.5058_*N*_0.2585_*P*_0.0229_*S*_0.0056_, which corresponds to 0.45 gC gDW^−1^ or 0.037 Cmol gDW^−1^, we obtain *D*_max_ = 132 kJ Cmol^−1^ h^−1^. To calculate biomass yield at a given growth rate, we separate the contributions from fermentation and respiration. For fermentation, Eqs. (5), (8), and (9) give:

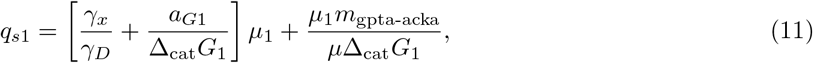

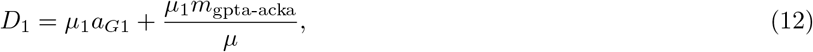

for pure respiration:

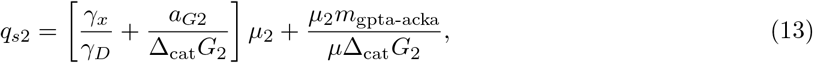

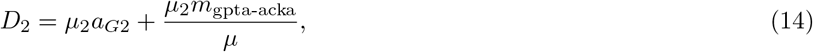

such that *q*_*s*_ = *q*_*s*1_ + *q*_*s*2_ and *D* = *D*_1_ + *D*_2_. Since fermentation requires less Gibbs energy per C-mol of biomass than respiration, it holds that:

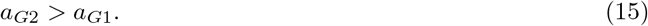

Moreover, we assume that:

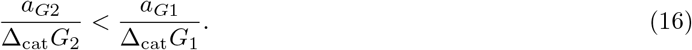

Which is clearly fulfilled for the numerical values mentioned in Table 1.

Before overflow metabolism, only pure respiration or fermentation is used(*µ* = *µ*_1_ *or µ* = *µ*_2_), according to the most economical strategy is pure respiration. For aerobic growth, overflow metabolism occurs at *µ >* (*D*_max_ − *m*_gpta-acka_)*/a*_*G*1_. At this growth rate, both *a*_*G*2_ and *a*_*G*1_ are much greater than *m*_gpta-acka_*/µ*. Therefore, the inequality assumption remains valid:

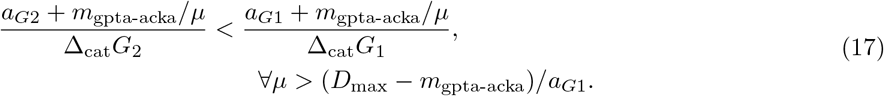

Which implies that, at fixed growth rate(*µ*), minimizing total carbon uptake is equivalent to maximizing the respiratory fraction according to (11), (13), (16) and (17). For growth rates

*µ < µ*_crit_ = (*D*_max_ − *m*_gpta-acka_)*/a*_*G*2_, the most economical strategy is therefore pure respiration. For growth rates *µ > µ*_max_ = (*D*_max_ − *m*_gpta-acka_)*/a*_*G*1_, no solution satisfies both (7) and (10).

For intermediate growth rates, *µ*_crit_ *< µ < µ*_crit_, the relative contributions of respiration and fermentation are obtained by solving (7) and (10). Introducing *c* = *a*_*G*2_ − *a*_*G*1_ and *r* = *D*_max_ − *m*_gpta-acka_ gives:

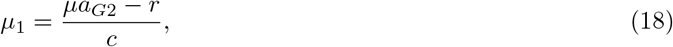

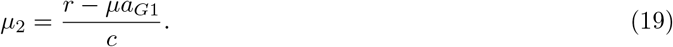

Given a specific carbon source at fixed growth rate, in order to calculate yield, we should first examine whether *µ > µ*_crit_. If *µ < µ*_crit_, the only respiration pathway is used(*µ* = *µ*_1_), yield can be derived through (11):

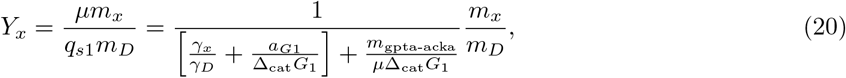

for *µ > µ*_crit_, respiration and fermentation pathways are both used, yield is generated from (18), (19), and (9):

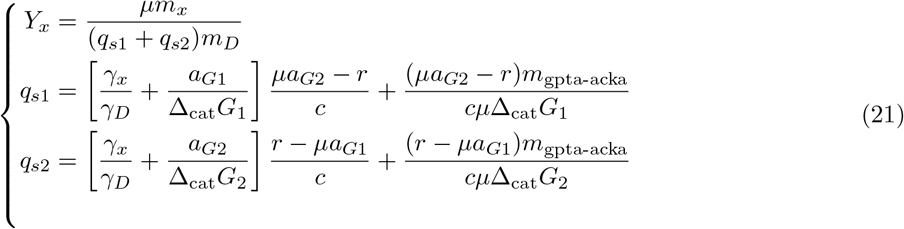

*µ* = *µ*_crit_ is regarded as the onset of overflow metabolism, beyond which the rate–yield relationship changes markedly owing to the recruitment of fermentative pathways(Fig. 1).

**Fig 1.**
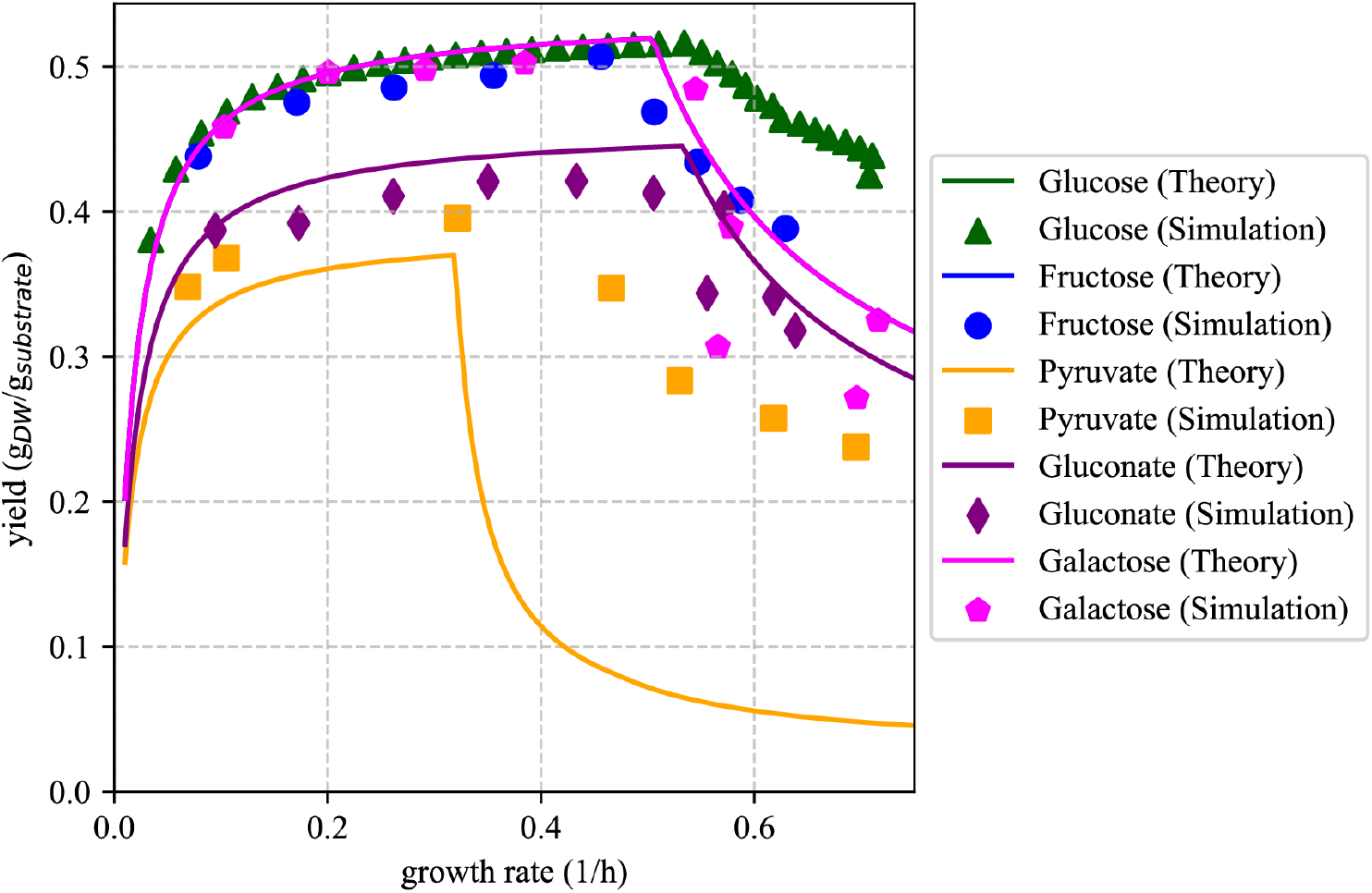
The relationship between Biomass yield and growth rate across five different substrates(glucose,fructose,pyruvate,gluconate,galactose). Solid lines indicate black box reconstruction using Heijen’s approximation parameters and symbols indicate genome scale thermoFBA simulation results.

Fig. 1 compares five substrates(glucose, fructose, galactose, pyruvate, and gluconate) using extended black box model with emperical parameters(see Table.S1,S3,S4) predictions (solid lines) and thermodynamic flux balance analysis (thermoFBA) simulation results (dots). As shown in the figure, the extended black box model successfully captures the transition to overflow metabolism, which is primarily reflected by the turning points in the predicted curves. Glucose, fructose, and galactose have identical elemental compositions and therefore share the same degree of reduction, carbon content, and catabolic Gibbs free energy change, leading to the same yield rate relationship. Pyruvate and gluconate show distinct behavior. Because gluconate is compositionally closer to glucose, its biomass yield is determined largely by the Gibbs free energy dissipated per unit substrate mass, Δ*G*_cat_*/m*_*s*_. Glucose dissipates more energy per gram (15.795 kJ g^−1^) than gluconate (13.161 kJ g^−1^), corresponding to higher carbon efficiency and higher biomass yield. Although pyruvate dissipates even less energy per gram (12.971 kJ g^−1^), its elemental composition differs more strongly from that of biomass, which increases the energetic cost of anabolic conversion and leads to a lower biomass yield than for gluconate.However, the theoretical predictions differs largely with simulation results, indicating potential improvement in parameter approximation.

### From genome scale metabolite models to black box growth models

The black box model contains four energetic parameters: *a*_*G*1_ and *a*_*G*2_, the Gibbs energy dissipated per C-mol biomass in the fermentative and respiratory branches, and *m*_gpta-acka_, the corresponding maintenance-related terms differ between the pre-overflow and overflow regimes. In classical black box analyses, these parameters are approximated from empirical correlations. Fig. 1 shows obvious overflow metabolism yet large deviation between black box model predictions with simulation results. Black box model provides a parsimonious and computationally efficient representation of input output relationships, typically involving only a small number of parameters. However, their simplified structure offers limited insight into the underlying metabolic mechanisms. In contrast, genome scale metabolic model incorporate detailed biochemical and genetic information, enabling mechanistic analysis of intracellular metabolism, but at the cost of substantially greater model complexity and computational demand. To exploit the complementary strengths of these two approaches, we establish a bridge between the black box and genome scale modeling framework. We estimate parameters in black box model directly from thermodynamically constrained genome scale flux solutions.

For each carbon source, thermoFBA provides a feasible flux distribution and the Gibbs energy dissipation rate of each active reaction. The total Gibbs energy dissipation rate is therefore:

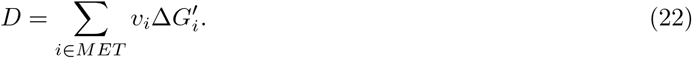

In the pure respiration regime (*µ*_1_ = 0), Eq. (8) becomes:

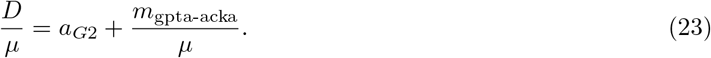

This predicts a linear relation between *D/µ* and 1*/µ*, with intercept *a*_*G*2_ and slope *m*_*g*_ as shown in Fig. 2.Remarkably, the same linear relationship emerges from genome scale model simulations, demonstrating a direct correspondence between the macroscopic thermodynamic description of the black box model and the detailed metabolic representation of the genome scale model. This agreement indicates that the thermodynamic constraints governing cellular growth are preserved across different levels of model abstraction. The genome scale model resolves the metabolic processes underlying these thermodynamic parameters, while the black box model summarizes their macroscopic consequences through a compact coarse grained description. This correspondence establishes a bridge between two models with fundamentally different levels of description and highlights the consistency of their underlying thermodynamic principles. That relation is not limited to glucose. Fructose, pyruvate, gluconate, and galactose all follow the expected linear trend(Fig. S1). In the overflow regime, Eq. (8) becomes:

**Fig 2.**
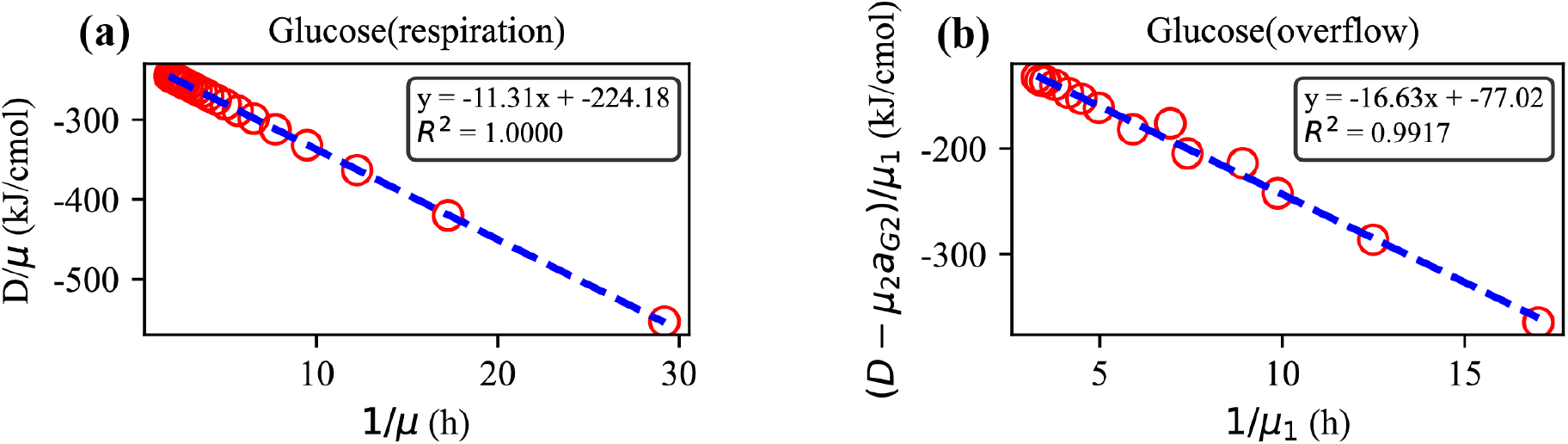
Linear relationships used to extract Gibbs energy dissipation parameters from the genome scale thermodynamic model for glucose. (a) Respiratory regime: plot of *D/µ* versus 1*/µ*, where the intercept gives *a*_*G*2_ and the slope gives *m*_gpta-acka_. (b) Overflow regime: plot of (*D* − *µ*_2_*a*_*G*2_)*/µ*_1_ versus 1*/µ*_1_, where the intercept gives *a*_*G*1_ and the slope gives *m*_gpta-acka_.

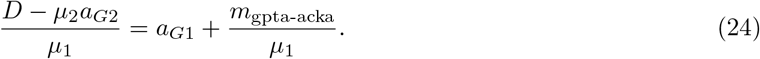

This predicts a second linear relation, with intercept *a*_*G*1_ and slope *m*_gpta-acka_.The results hold not only for glucose(Fig. 2) but also for fructose, pyruvate, gluconate, and galactose(Fig. S2).For glucose, fructose, and pyruvate, a clear linear relationship is observed, whereas galactose and gluconate retain an approximately linear trend with larger deviations. One possible explanation is the formation of additional metabolic byproducts during the metabolism of these two carbon sources, which would violate the model assumption that metabolism can be represented by only two metabolic pathways. Thus, the expected black box linear relationships are recovered for both respiration and overflow growth across multiple carbon sources. This indicates that the black box model and the genome scale model are consistent at the level of Gibbs energy dissipation, and therefore provides the thermodynamic basis for linking these two descriptions.

The extracted parameters are summarized in Table. S1 and Table. S2, and compared with the corresponding empirical estimates in Fig. 3. The closest agreement is found for the respiratory dissipation coefficient *a*_*G*2_ (Fig. 3b) and *m*_gpta-acka_(Fig. 3d) after overflow. By contrast, *a*_*G*1_ and *m*_gpta-acka_ before overflow extracted from genome scale model are generally lower than those predicted by Heijnen’s approximation(Fig. 3a,c). This difference is expected. Heijen’s estimation are coarse grained and depend only on a limited set of substrate properties: degree of reduction, carbon number and temperature.For instance, glucose, fructose, and galactose have the same degree of reduction and the same number of carbon atoms, so Heijen’s relation predicts the same *a*_*G*_. The genome scale model instead resolves pathway details, and therefore yields differences across these substrates that the Heijnen’s relation cannot capture. More interestingly, *m*_*g*_ is generally higher under overflow conditions than before the onset of overflow metabolism. This information cannot be obtained directly from Heijnen’s empirical correlation. The elevated maintenance requirement may reflect the simultaneous operation of respiration and fermentation during overflow metabolism, which increases the number of enzyme catalyzed reactions that must be maintained. Consequently, the greater demand for enzyme turnover, repair, and replacement may contribute to the higher maintenance energy dissipation observed under overflow conditions.

**Fig 3.**
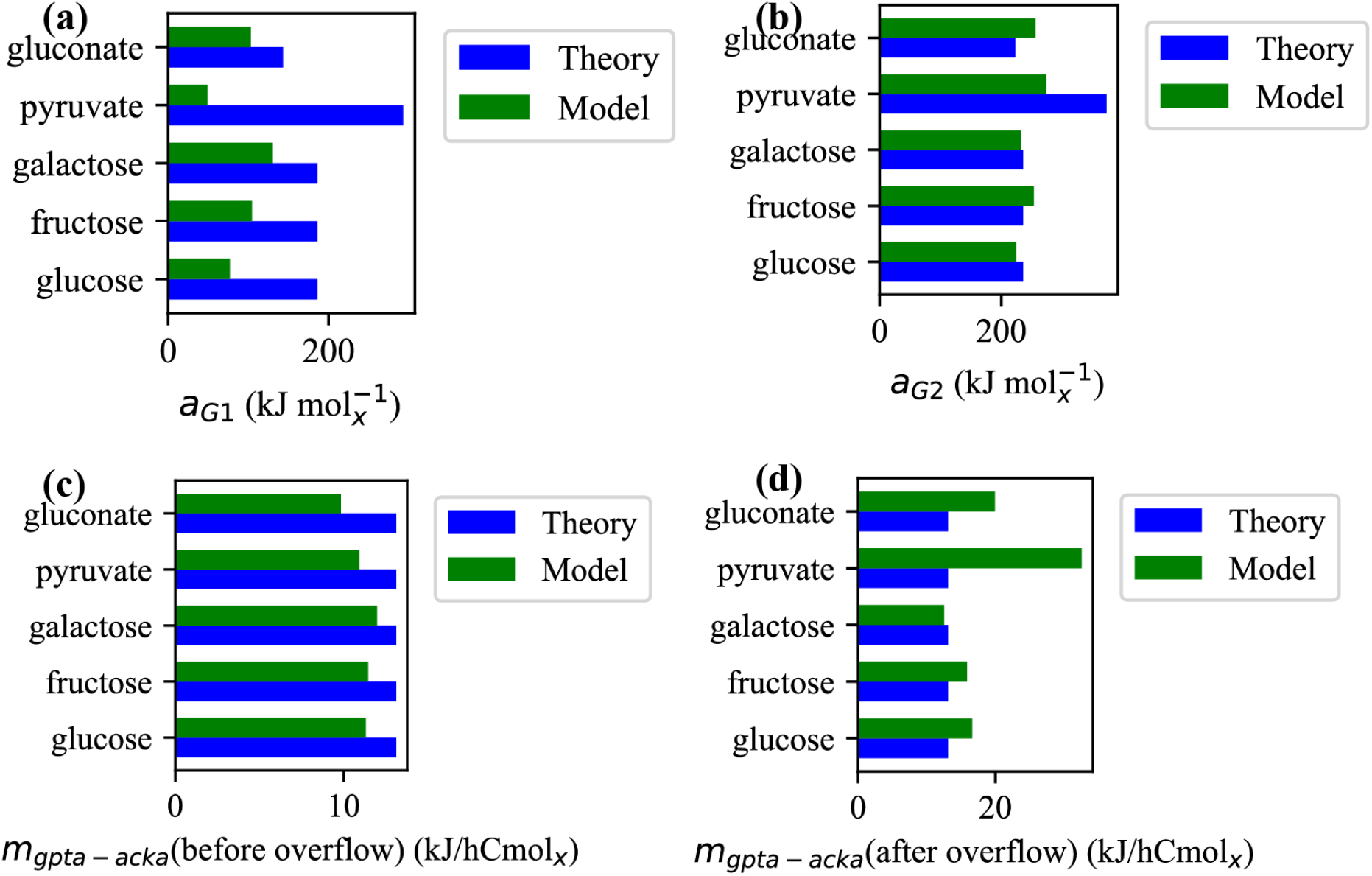
Comparison of energetic parameters (a) fermentation Gibbs energy dissipation rate per C-mol biomass formed*a*_*G*1_, (b) respiration Gibbs energy dissipation rate per C-mol biomass formed *a*_*G*2_, (c) respiration maintenance energy *m*_gpta-acka_, and (d) overflow maintenance energy *m*_gpta-acka_ for glucose, fructose, galactose, pyruvate, and gluconate. Blue bars denote Heijen’s approximation relation, green bars denote values extracted from thermodynamically constrained genome scale model.

In Fig. 4a, we examine whether the genome scale derived black box parameters are sufficient to reconstruct experimentally observed bacterial physiology. The predicted growth curve (black line) shows close agreement with experimental data (red circles and triangles), indicating that the simplified model successfully preserves the quantitative relationship between substrate uptake and cellular growth. Beyond growth prediction, the model naturally recovers the characteristic signatures of overflow metabolism. At a growth rate of approximately 0.5 h^−1^, acetate secretion emerges as the respiratory capacity becomes insufficient to support further biomass production, leading to the activation of fermentative metabolism. As a consequence, oxygen uptake and carbon dioxide production decrease after the metabolic transition, consistent with the experimentally observed shift from respiration dominated growth to mixed respiratory and fermentative growth. Thus, the black box model captures not only the growth rate but also the underlying physiological transition associated with overflow metabolism.

**Fig 4.**
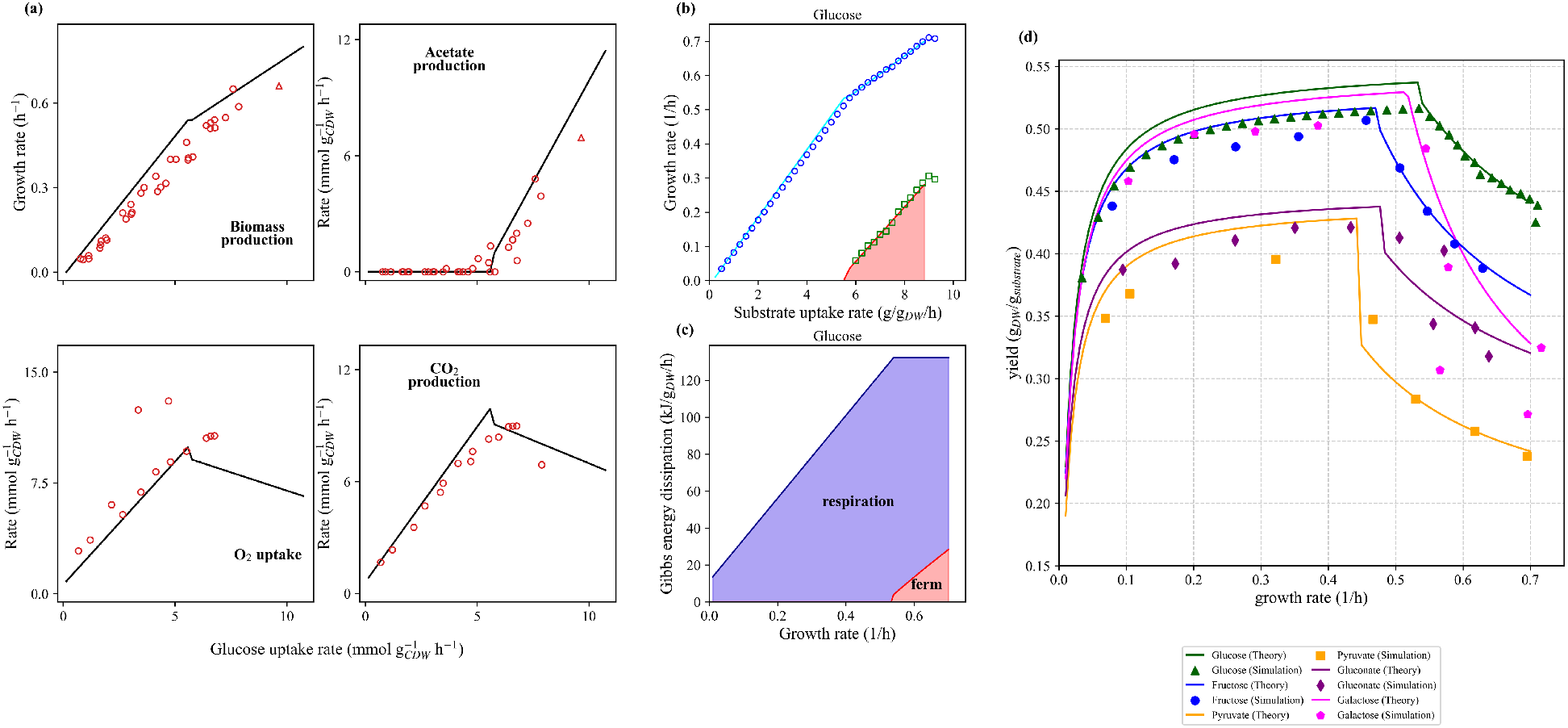
Reconstruction of black box model using parameters extracted from genome scale thermodynamic model. (a)Black box growth model prediction(black lines) fit well with experimental data in biomass production, acetate excretion, *O*_2_ uptake rate and *CO*_2_ production rate.Red circles represent experimentally determined values from glucose-limited chemostat cultures [39–43], and red triangles values from glucose batch culture [44] (b) The relationship between growth rate and substrate uptake rate. Blue line denote growth rate predicted by black box model using extracted parameters and the red line denote fermentation growth rate predicted by black box model using extracted parameters. Symbols indicate simulation derived values, in which blue represent total growth rate and green represnet fermentation part. (c) The relationship between Gibbs energy dissipation rate and growth rate. Red branch denote fermention while blue branch denote respiration. (d) The relationship between Biomass yield and growth rate across five different substrates(glucose,fructose,pyruvate,gluconate,galactose). Solid lines indicate black box reconstruction using extracted parameters and symbols indicate genome scale thermoFBA simulation results.

To further evaluate the consistency between the two modeling approaches, we directly compare the black box model predictions with the corresponding genome scale model simulations. In Fig. 4b, the blue points represent the genome scale relationship between growth rate and glucose uptake rate. The blue line shows the corresponding black box prediction obtained using genome scale derived parameters together with the overflow extension, and it closely reproduces the genome scale simulation results. The green points represent the respiratory contribution to growth calculated from the genome scale model, while the green line shows the corresponding black box prediction. These results also exhibit excellent agreement, demonstrating that the black box model successfully captures the partitioning between respiratory and fermentative growth. Similar consistency is observed for other carbon sources in Fig. S3a1–d1.

Fig. 4c shows the partition of Gibbs energy dissipation during glucose growth calculated by the black box model using the extracted parameters and the overflow extension. Such a separation of energy dissipation between respiratory and fermentative metabolism is difficult to obtain directly from genome scale models, because respiration and fermentation share common upstream reactions, such as the glycolytic pathway, and their contributions cannot be uniquely assigned without further defining the downstream reactions as respiratory or fermentative processes. In contrast, the black box model naturally separates the two metabolic modes according to their distinct thermodynamic contributions. The blue region represents respiratory dissipation, whereas the red region represents fermentative dissipation. Fermentative dissipation emerges only after the onset of overflow metabolism. Even at the maximal growth rate, fermentation contributes only approximately one quarter of the total dissipation, indicating that *E. coli* maintains substantial respiratory activity after overflow begins. The same pattern is observed for other carbon sources in Fig. S3a2–d2.

Fig. 4d shows the rate yield relation for all carbon sources. Filled symbols are genome scale simulations and solid lines are black box predictions obtained using the extracted parameters and specific extension. For all substrates, the curves rise and then decline because overflow begins beyond a critical growth rate and cells shift from carbon efficient respiration to less efficient but faster ATP producing fermentation. The close agreement between theory and simulation indicates that genome scale behavior can be accurately reconstructed by a extended black box model with a finite set of thermodynamic parameters.

These results show that Gibbs energy dissipation parameters do not need to be treated as purely empirical fitting constants external to genome scale metabolism. Instead, they can be extracted directly from genome scale thermoFBA solutions, interpreted through the respiratory and overflow branches of the black box model, and then used to reconstruct the macroscopic rate yield behavior. In this way, the genome scale model provides the mechanistic basis of the black box parameters, whereas the black box model provides a compact thermodynamic summary of the genome scale model’s solution behavior. This correspondence further supports the interpretation of the black box model as an effective coarse grained representation of the genome scale model. Because genome scale models are formulated from stoichiometrically balanced biochemical reactions subject to linear mass conservation constraints, their fundamental structure is consistent with the linear assumptions underlying the black box formulation. Importantly, the reconstructed black box predictions also agree with experimentally observed bacterial physiology, including growth behavior and the emergence of overflow metabolism.

### Extending the thermodynamic connection to yeast

To further examine the generality of the proposed thermodynamic connection across organisms, we applied the same procedure to a thermodynamically constrained genome scale model of *S. cerevisiae*. The model integrates a stoichiometric metabolic network with thermodynamic constraints, including mass balance, proton and charge balance, Gibbs energy balance, and the second law of thermodynamics. It describes 241 metabolic processes and 156 metabolites, providing a detailed representation of cellular metabolism while maintaining thermodynamic consistency.

Following the workflow established for *E. coli*, we extracted the effective Gibbs energy dissipation parameters of the black box model from genome scale simulations. As shown in Fig. 5, the extracted parameters again satisfy the predicted linear relationship between *D* and *µ*, demonstrating that the coarse grained thermodynamic description emerges naturally from the genome scale model. Interestingly, the extracted growth associated dissipation parameter is *m*_*g*_ = 0, which is consistent with the maintenance energy setting of the yeast genome scale model. This agreement further confirms that the effective parameters of the black box model preserve the thermodynamic constraints encoded in the genome scale model.

**Fig 5.**
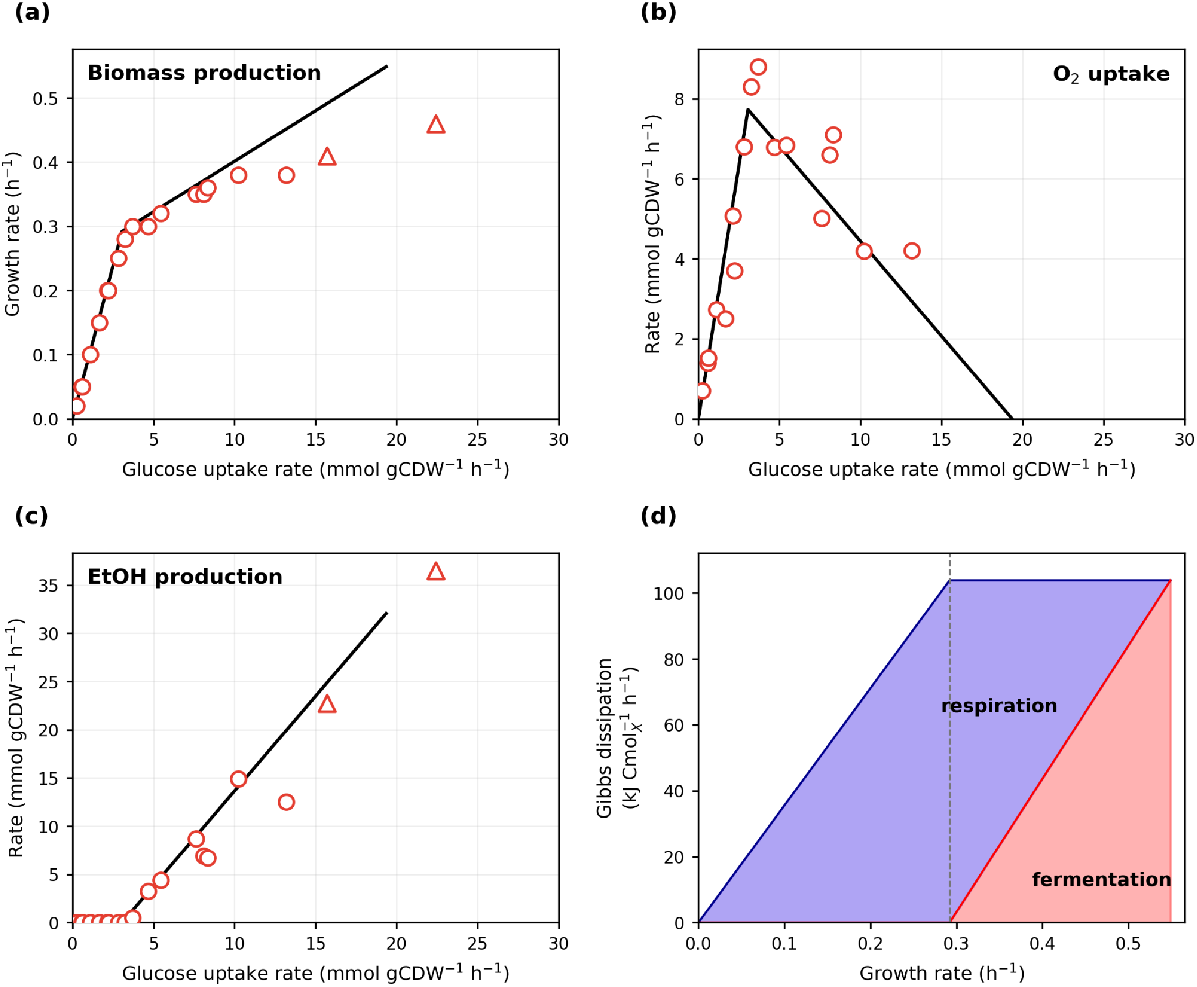
Reconstruction of the yeast black box model using parameters extracted from genome scale thermodynamic model. (a–c) Comparison between black box predictions and experimental measurements for biomass production, oxygen uptake, and ethanol production.Red circles represent experimentally determined values from glucose-limited chemostat cultures [45, 46] and red triangles values from glucose batch cultures [46, 47] (d) Gibbs energy dissipation partition predicted by the black box model. Unlike *E. coli*, fermentation contributes nearly all dissipation at the maximal growth rate.

Using these extracted parameters, the black box growth model was then applied to predict yeast physiology. As shown in Fig. 5, the predicted growth, oxygen uptake, and ethanol production rates agree well with experimental measurements over a wide range of glucose uptake rates. The model also reproduces the transition from respiration to fermentation at increasing substrate uptake rates. Analysis of the corresponding Gibbs energy dissipation pathways reveals a distinct metabolic strategy compared with *E. coli* : at the maximum growth rate, fermentation accounts for nearly all Gibbs energy dissipation, whereas respiratory dissipation remains significant in *E. coli*.

These results demonstrate that the connection between genome scale thermodynamic models and black box growth models is not restricted to a specific organism. Instead, once a genome scale model is constructed with appropriate thermodynamic constraints, its complex metabolic behavior can be systematically coarse grained into a small number of effective thermodynamic parameters. Therefore, the computationally efficient black box model and the detailed but computationally demanding genome scale model can describe cellular physiology within a unified thermodynamic framework.

## Discussion

In this work, we establish a quantitative connection between black box models of microbial growth and genome scale thermoFBA (thermodynamic flux balance analysis) frameworks. Black box models describe energetic limits of growth using macroscopic parameters such as Gibbs free energy dissipation and maintenance energy, but they do not explicitly resolve the intracellular processes underlying these relationships. Genome scale metabolic models, by contrast, represent metabolism at the level of individual reactions and pathways, but they do not directly provide the macroscopic energetic parameters used in black box models. By exploiting the linear relations between genome scale thermoFBA solutions and black box model variables, we establish a thermodynamic bridge between black box growth models and genome scale models for both *E. coli* and *S. cerevisiae*.The agreement among genome scale model predictions, reconstructed black box model predictions, and experimental physiological data demonstrates that genome scale metabolic information can be systematically transferred into a low dimensional thermodynamic description while preserving the experimentally observed cellular physiology.Therefore, this thermodynamic connection provides a general strategy for linking black box growth models with genome scale FBA models constructed with consistent thermodynamic constraints across different organisms.

thermodynamic flux balance analysis (thermoFBA) predicts metabolic states through genome scale models, generating flux solutions that encode high dimensional information arising from complex metabolic networks. In contrast, the black box model describes microbial growth using a small number of thermodynamic parameters. Here, we demonstrate that genome scale thermoFBA models and black box model are thermodynamically consistent, showing that the high dimensional information contained in thermoFBA solutions can be effectively captured by a low dimensional thermodynamic representation while retaining key growth properties.

While the present framework captures the thermodynamic consistency between genome scale models and black box descriptions, several aspects remain open for future investigation. The analysis depends on thermodynamic parameters extracted from a specific genome scale reconstruction, and therefore its accuracy is influenced by the quality of the underlying metabolic network and thermodynamic data. In addition, the black box formulation represents microbial growth through simplified respiratory and fermentative states, while actual metabolism may involve more diverse and interconnected pathway combinations. Furthermore, regulatory mechanisms, enzyme allocation constraints, and kinetic effects are not explicitly captured, although these factors can shape both metabolic flux distributions and Gibbs energy dissipation in living systems. Future integration of thermodynamic modeling with enzyme constrained or resource allocation models may provide a more comprehensive quantitative description of microbial physiology.

Overall, these results show that classical black box models and modern genome scale metabolic models are not competing descriptions, but complementary views of the same biological system. By linking macroscopic energetic parameters to intracellular metabolic fluxes, this study provides a multiscale thermodynamic framework for understanding microbial metabolism and a basis for integrating energetic theory with genome scale systems biology.

## Appendix Construction of thermodynamic flux balance analysis

thermodynamic flux balance analysis (thermoFBA), proposed by Niebel et al. [27], extends classical flux balance analysis by incorporating thermodynamic constraints into metabolic network analysis. Their study demonstrated that cellular metabolism is subject to an upper bound on Gibbs free energy dissipation. By imposing this constraint, the model successfully predicts cellular physiology and intracellular flux distributions across different glucose uptake rates, as well as the maximal achievable growth rate.

Similar to conventional FBA, the model assumes steady state mass balance for intracellular metabolites,

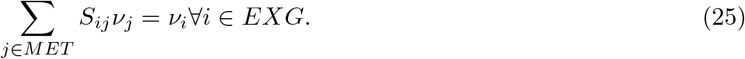

In addition to mass balance, thermoFBA introduces constraints on cellular Gibbs energy dissipation. The total Gibbs energy dissipation rate *D* is defined as the sum of Gibbs energy exchange rates across the system boundary and the internal Gibbs energy dissipation rates within the metabolic network,

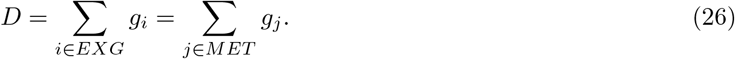

The Gibbs energy exchange rate associated with exchange reactions is defined as

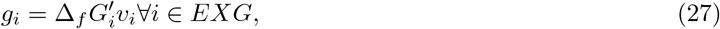

where 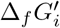 denotes the Gibbs free energy of formation of metabolite *i*.

The Gibbs energy dissipation rate of intracellular metabolic reactions is given by

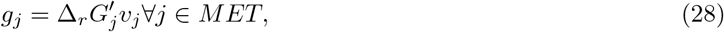

where 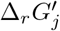 represents the Gibbs free energy change of metabolic reaction *j*. The reaction Gibbs free energy is determined by

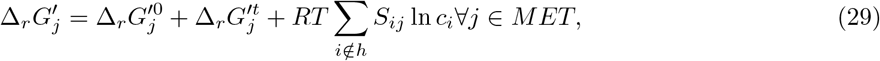

where 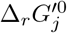 denotes the standard Gibbs free energy of reaction, 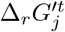 accounts for Gibbs energy contributions associated with metabolite transport, *c*_*i*_ is the intracellular concentration of metabolite *i, T* is the absolute temperature, and *R* is the universal gas constant.

The Gibbs free energy of formation of exchanged metabolites is calculated as

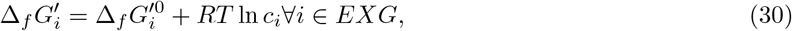

where 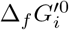 denotes the standard Gibbs free energy of formation of metabolite *i*.

Although metabolic reactions are assumed to be reversible in principle, feasible flux directions must satisfy the second law of thermodynamics. Therefore,

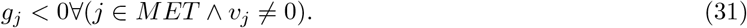

Combining the stoichiometric and thermodynamic relations above yields the thermodynamic stoichiometric model

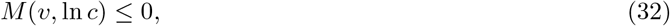

which defines the feasible space of flux distributions *v* and metabolite concentrations ln *c* satisfying both mass balance and thermodynamic constraints:

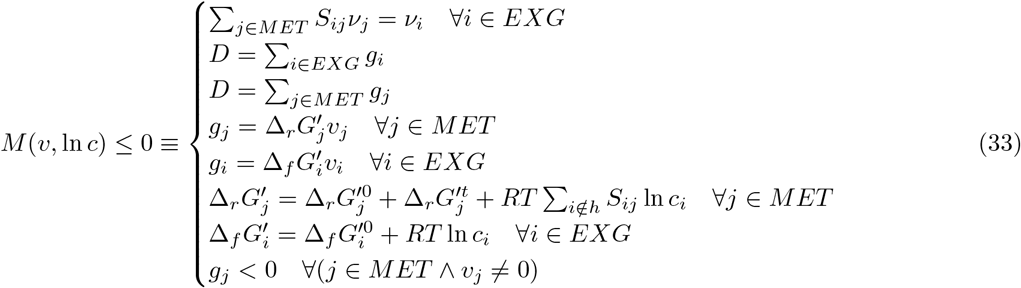

## Supporting information

**S1 Table**. Comparison of theoretical and genome-scale extracted Gibbs energy dissipation parameters (see the accompanying supporting-information source).

**S2 Table**. Comparison of theoretical and genome-scale extracted maintenance-energy parameters.

**S3 Table**. Summary of parameters used in the black-box model without genome-scale information.

**S4 Table**. Substrate-dependent parameters calculated from molecular formulae.

**S1 Fig**. Respiration-associated energetic-parameter extraction for non-glucose substrates.

**S2 Fig**. Fermentation-associated energetic-parameter extraction for non-glucose substrates.

**S3 Fig**. Reconstruction of the black-box model across non-glucose substrates.

**S4 Fig**. Extraction of Gibbs energy dissipation parameters for yeast.

## Acknowledgments

This work was supported by the National Natural Science Foundation of China(Grants No.12274426 and No.12447101). The authors have declared that no competing interests exist. Pujiang Jia gratefully acknowledges Matthias and Edward N. Smith for helpful discussions by email that aided his understanding of the genome scale model. Useful discussions with Siyu Li, Maggie Wang, Liang Zhang, Shaohua Guan, Zhichao Zhang are gratefully acknowledged.

## Data Availability

The data supporting the findings of this study are generated from numerical simulations. The thermoFBA source codes and model implementations for both *E. coli* and *S. cerevisiae*, together with the parameters and simulation settings required to reproduce the reported results, are publicly available in Zenodo (DOI: 10.5281/zenodo.21849804). The experimental data used for model comparison and validation were obtained from previously published studies cited in the manuscript.

